# *Candidatus (Ca.)* phytoplasma asteris subgroups display distinct disease progression dynamics during the carrot growing season

**DOI:** 10.1101/2020.09.17.301150

**Authors:** Justin Clements, Benjamin Z. Bradford, Marjorie Garcia, Shannon Piper, Weijie Huang, Agnieszka Zwolinska, Kurt Lamour, Saskia Hogenhout, Russell L. Groves

## Abstract

Aster Yellows phytoplasma (AYp; *Candidatus (Ca.)* Phytoplasma asteris) is an obligate bacterial pathogen that is the causative agent of multiple diseases in herbaceous plants. While this phytoplasma has been examined in depth for its disease characteristics, knowledge about the spatial and temporal dynamics of pathogen spread is lacking. The phytoplasma is found in plant’s phloem and is vectored by leafhoppers (Cicadellidae: Hemiptera), including the aster leafhopper, *Macrosteles quadrilineatus* Forbes. The aster leafhopper is a migratory insect pest that overwinters in the southern United States, and historical data suggest these insects migrate from southern overwintering locations to northern latitudes annually, transmitting and driving phytoplasma infection rates as they migrate. A more in-depth understanding of the spatial, temporal and genetic determinants of Aster Yellows disease progress will lead to better integrated pest management strategies for Aster Yellows disease control. Carrot, *Daucus carota* L., plots were established at two planting densities in central Wisconsin and monitored during the 2018 growing season for Aster Yellows disease progression. Symptomatic carrots were sampled and assayed for the presence of the Aster Yellows phytoplasma. Aster Yellows disease progression was determined to be significantly associated with calendar date, crop density, location within the field, and phytoplasma subgroup.

## Introduction

Understanding the progression of pathogen infections resulting in disease phenotypes in agricultural crops is critically important for determining effective pest management strategies. Aster Yellows phytoplasma (AYp), *Candidatus* (*Ca*.) Phytoplasma asteris is one agricultural pathogen of concern, particularly for commercial growers of carrot, lettuce and celery in the Midwest. This small, wall-less prokaryote can infect more than 350 different species of plants including vital agricultural crops resulting in multiple host-dependent disease phenotypes [1-3]. In carrot, the AYp pathogen causes Aster Yellows (AY) disease, which is characterized by stunted growth, bittering of the root, and premature plant death. This pathogen is vectored by multiple leafhopper species [1-3]. However, in the central United States, the primary vector of concern is the aster leafhopper (ALH), *Macrosteles quadrilineatus* Forbes (Hemiptera: Cicadellidae), which enables the movement of the phytoplasma from infected hosts into susceptible healthy plants. ALH is a migratory insect, overwintering in the southern United States [3]. Historically, the migration of ALH from the southern United States into northern, agriculturally-intensive areas is thought to drive infection rates (number of AYp infected leafhoppers/ total number of leafhoppers collected) [1].

Understanding and reducing the spread of pathogenic agents in the field is of critical importance for agricultural growers. The movement of AYp across larger, geographic areas has previously been examined in depth [1-3]. However, the progression and movement of AYp genotypes within individual commercial agricultural fields has not been explored. Previous studies have examined different disease models, and determined several factors influencing the movement of pathogens within different plant species [4-7]. These include insect sensory mechanisms [7], pathogen-protein manipulation of host plants [8], the movement of pathogens within the host plant [4], and geographic and temporal components [3,9]. These results suggest multiple factors are involved in the development of endemic disease conditions resulting in AY. Characterizing the specific factors, including the arrival and colonization of carrot fields by the ALH, and the development and spread of the AY disease vectored by these leafhoppers, would provide growers with additional information useful for managing this economically important disease.

Symptoms of AY disease can vary depending on the infected plant species. In commercial carrots, *Daucus carota* L., AYp can cause the agriculturally significant disease AY. Symptoms include witch’s broom (proliferation of shoots), phyllody (retrograde development of flowers into leaves), virescence (flower organs remaining green), bolting, formation of shortened internode and elongated petioles, chlorosis, and the production of hairy roots [10]. The etiology of Aster Yellows related disease consists of one taxon group of phytoplasma, *Candidatus* (*Ca*.) Phytoplasma asteris. However, within the species designation there are multiple subgroups of the phytoplasma [8, 11-12]. The subgroup classification is based on the phylogenetic difference between a conserved region of the 16S ribosomal RNA [13]. Within the Midwestern region of the United States, and more specifically Wisconsin, the AYp subgroups 16SrI-A and 16SrI-B have been detected and are both known causative agents of AY in carrots [2].

In the 16SrI-A AYp strain, secreted AY-WB protein (SAP) effector genes have been identified and some have been shown to promote phytoplasma virulence and enhanced insect-pathogen and insect-plant interactions [14-16]. Studies have identified 56 putative SAPs in the AY-WB genome and SAP11 was found to migrate out of the phloem into adjacent tissues, including trichomes, of *A*. *thaliana* [17]. Secreted AY-WB11 is a known, destabilizing class II TCP transcription factor of *Arabidopsis thaliana* leading to the induction of shoot branching resembling witch’s broom symptoms and leaf crinkling [16, 18-22]. Moreover, SAP11 suppresses jasmonic acid (JA) synthesis and modulates volatile organic compounds (VOC) originating from glandular trichomes of plants [16,21]. This interaction relies on a nuclear localization signal in SAP11, which has also been predicted in SAP22, SAP30 and SAP42 [17]. The function of another effector, SAP54, has also been characterized in depth, and is known to result in phenotypic changes in leaf-like sepals, fewer and less elongated siliques, loss of floral determinacy, and other reproductive deficiencies [14]. Secreted AY-WB 54 interacts with type II MADS-domain transcription factors (MTFs) and the 26S proteasome shuttle factor RAD23 leading to degradation of MTFs in plants, causing the phyllody phenotype [14]. The SAP54 effector is also responsible for enhanced colonization by insect vectors on plants [15], a phenotype that is dependent on SAP54 interaction with RAD23 [15]. Tomkins et al. used a multi-layered model to predict the complex interactions between leafhopper, phytoplasma and plants related to the expression of effector genes SAP11 and SAP54 and suggested that the effectors contribute to disease spread [6]. The majority of SAP genes lie in close proximity to each other on apparent pathogenicity islands that resemble conjugative transposable elements that were named potential mobile units (PMUs) [22-23]. These PMUs are variably present and horizontally exchanged among phytoplasmas [24]. In the genome of AY-WB phytoplasma, the gene for SAP11 lies on a PMU-like genomic region adjacent to other candidate effector genes encoding for SAP56, SAP66, SAP67, SAP68. The genes for 34 of the 56 SAPs are also located on PMUs, while 7 are located on plasmids [17, 20, 22].

In the current investigation, our overarching goal was to investigate the disease progression of AY within a carrot field, emphasizing the spatial and temporal dynamics of AY incidence and the composition of SAP effectors genes associated with AYp strains found in infected plants. We further examined the effects of within-field location, time, and planting density on AY incidence and characterized the genetic markers related to the composition of AYp. We found that the patterns of disease spread are not random but structured, and are the result of multiple biological parameters. The findings provide insight into the movement and colonization of AYp in a carrot field and will lead to a better understanding of disease progression and management.

## Materials and Methods

## Data availability

All relevant data are contained within the paper and its supporting information files.

## Ethical statement

This article does not contain studies with any human participants and no specific permits were required for field collection or experimental treatment of *Macrosteles quadrilineatus* for the study described.

### Disease progression and leafhopper pressure

On May 7, 2018, carrot plantings (cv. ‘Canada’) were established at the University of Wisconsin’s Hancock Agricultural Research Station (HARS) in field K3 (44.118181°N, -89.549045°W). Seed was purchased through a commercial vendor (Seedway LLC) which conducts phytosanitary procedures necessary to screen for pathogens and was certified disease free. Within a larger, 180m by 45m carrot field, a central 66m wide section was divided in half and planted at either a high density (1360k seeds per hectare) or low density (556k per hectare) seeding rate. Each planting was further divided into five rows of 18 beds with 4m bare alleys separating rows (90 beds per density). Each 3-row raised bed measured 2m wide by 6m long. The first and last rows of beds were considered “edge” beds and were bordered by other, non-carrot crops to the north (sorghum) and the south (green bean). Carrot beds were subsequently managed with a standard fertilizer program, but no crop protection pesticides (herbicides, fungicides, insecticides) were applied at any point in the season. Weeds were managed by hand-pulling every two weeks.

Using a random number generator, two beds from each row of the high- and low-density plantings were selected as plots for biweekly stand counts and AY disease monitoring. Initial counts occurred on Jun 27, 2018, when plots were staked, stand counts of all carrots within selected beds were conducted, and AY disease phenotype of each carrot observed. Possible AY disease phenotypes included proliferation of shoots (“witches broom”), retrograde development of flowers into leaves, virescence, formation of shortened internode and elongated leaves, and yellowing/reddening of foliage. Plots were scouted in this way every two weeks (Jun 27, Jul 11 and 25, Aug 8 and 21, Sep 6 and 18, Oct 2 and 16) for a total of 9 observations. The exact location of each phenotypically-described, diseased carrot was recorded and marked within the field relative to the front stake of each row. Each symptomatic carrot had a small sample of both petiole and stem removed for genetic confirmation of the presence of the AYp pathogen in the carrot tissue. Symptomatic carrots were tagged with a small plastic stake to prevent double-counting over successive sample dates.

To assess ALH abundance and infection prevalence within the leafhoppers within the carrot fields, 1000 sweeps of the plant canopy were performed in both the high- and low-density plantings using a standard, 15-inch sweep net during each biweekly disease monitoring event. Collected insects were bagged in 3.8 L sealable plastic bags and placed on ice for transportation to the University of Wisconsin-Madison for processing, where the total number of ALH was determined and a subset of individuals (N=40) obtained from each planting density/sample time were stored for later analysis. A taxonomic key to the family Cicadellidae was used to confirm the identity of all adult ALH [25].

### DNA isolation from plant and leafhopper tissue

Tissue samples from all symptomatic carrots and 40 leafhoppers per density/sample time were analyzed. Tissue samples (10 mg) collected from carrots were processed as individual samples based upon tissue type (petiole, stem). Samples were placed in corresponding 1.5 ml sterile microcentrifuge tubes and homogenized with a sterile plastic pestle in 500 µl 2% CTAB (Cetyl-trimethyl-ammonium-bromide) (bioWORLD, Dublin, OH) buffer with 1 µl of 0.2ng/µl RNase A (Thermo Fisher Scientific, Waltham, MA). Homogenates were incubated at 60°C for 30 min, centrifuged at 12,000g for 5 minutes, and the supernatant was transferred to a fresh tube. A single volume of chloroform was added and the samples were gently mixed for 10 minutes. Samples were then centrifuged for 10 minutes at 12,000g and the supernatant was again transferred to a fresh tube. DNA was precipitated by adding 1 volume of cold isopropanol and mixed by inversion for 10 minutes. Samples were centrifuged to pelletize DNA and washed with 75% EtOH. After washing, EtOH was discarded and samples were allowed to completely air dry; methodology was adapted from Marzachi *et al*. [26]. DNA was suspended in DNase/ RNase free H_2_O, and subsequently quantified using a NanoDrop, microvolume spectrophotometer (Thermo Fisher Scientific, Waltham, MA), and brought to a final volume of between 20 µl -100 µl depending on the measured concentration. Samples were then frozen at -20°C for further analysis.

### Confirmation of Aster Yellows phytoplasma within tissue samples

To confirm the presence of AYp in DNA isolations from tissue samples, a P1/P7 PCR amplification was conducted, followed by a R16F2n/R16R2 nested PCR on the P1/P7 PCR product to amplify the 16S rRNA sequence. DNA primers were obtained from Smart *et al*. 1996 (P1/P7) [27] and Gundersen *et al*. 1996 (R16F2n/R16R2) [28] and are presented in **Supplementary Table S1**. Specifically, 25 µl PCR reactions were conducted with GoTaq® Green Master Mix (Promega Corporation, Madison, WI). Reaction conditions included 240 seconds at 94°C as the initial denaturing step, followed by 30 cycles of 30 seconds at 94°C for denaturation, 60 seconds at 64°C (P1/P7) or 60°C (R16F2n/R16R2) for annealing, 90 seconds at 72°C for extension and a final extension of 300 seconds at 72°C. PCR amplification products were run on a 1.5% agarose gel to confirm the presence of corresponding DNA fragments. The corresponding DNA fragment observed from the P1/P7 amplification was 1.8 kbp, while the nested product was 1.2 kbp. Total DNA of only extracted carrot tissue through CTAB procedure was used to analyze the genetic composition of subgroup and effector proportions. Subgroup identification was determined using both nucleic acid sequencing and restriction fragment length polymorphism (RFLP). The identity and position of six unique, single nucleic acid polymorphisms (SNPs) were compared between sequences to identify AYp subgroup. Furthermore, a RFLP assay using the restriction endonuclease Hha1 (Promega Corporation), was conducted at 37°C for 90 min and run on 1.5% agarose gel. RFLP was used as a supplementary assay to confirm the subgroup designation resulting from SNP assessment of the sequencing data [11].

### Effector determination by ‘MonsterPlex’ sequencing

To assess the genetic variability of Aster Yellows phytoplasma approximately 400ng of each AYp-positive carrot sample’s DNA was submitted to Floodlight Genomics LLC (Knoxville, TN) for ‘MonsterPlex’ amplification and Illumina DNA sequencing. Floodlight Genomics used an optimized Hi-Plex approach to amplify targets in a single multiplex reaction with 21 targets including 16S rRNA and 20 effector sequences (**Supplementary table S2**). The sample-specific barcoded amplicons were sequenced on the Illumina HiSeq X platform according to the manufacturer’s directions. Floodlight Genomics delivered sample-specific raw DNA sequence reads as FASTQ files. Annotation of the raw reads was performed with Geneious bioinformatics software (Auckland, New Zealand). Raw reads were aligned to reference sequences and annotated. To determine subgroup designation, the reads were annotated for known SNPs associated with each subgroup [11]. Effector reads were aligned to reference sequences, and the total number of effector reads were standardized to the number of 16S rRNA within each sample to generate a reads ratio indicating the number of reads of each effector per 16S rRNA read. Effectors were classified as being present in a sample if the read number after standardization was greater than 1. Effectors SAP21, 36, 54, and 67 did not amplify in either subgroup and were removed from further analysis, as amplification might have been the result of primer efficiency within the ‘MonsterPlex’ analysis. Samples were considered AYp-infected if both the PCR and genetic sequencing were positive for the presence of the phytoplasma (16S rRNA sequences were searched against the non-redundant nucleotide NCBI database using BLASTn).

### Data analysis

The effect of date, planting density, and plot location (edge/interior of field) on AY disease incidence was quantified using a maximum likelihood regression model with a beta binomial distribution and a logit link function in JMP Pro 13.2.1 (SAS Institute, Cary, NC). Beta binomial models require a total count and a diseased count variable, both of which were generated for each plot on each scouting date. Differences in effector copy number by AYp subgroup were analyzed in R version 3.6.1 (R Core Team, Vienna, Austria) by running a binomial regression with a logit link function for each effector and performing a Chi-squared test on the resultant model.

## Results

### Edge- and density-dependent disease progression

Aster Yellows disease progression increased over the growing season and was also dependent upon initial planting density. On the first sampling date (27 Jun 2018), no symptomatic carrots were identified. Disease incidence gradually increased throughout the season until October, when it increased exponentially, reaching an average of 4.5 ± 1.8% (edge plots) and 4.3 ± 1.9% (interior plots) in the high-density planting, and an average of 11.0 ± 5.9% (edge plots) and 8.2 ± 2.2% (interior plots) in the low-density planting (**Figure 1, Supplemental Table S3)**.

**Figure 1.**
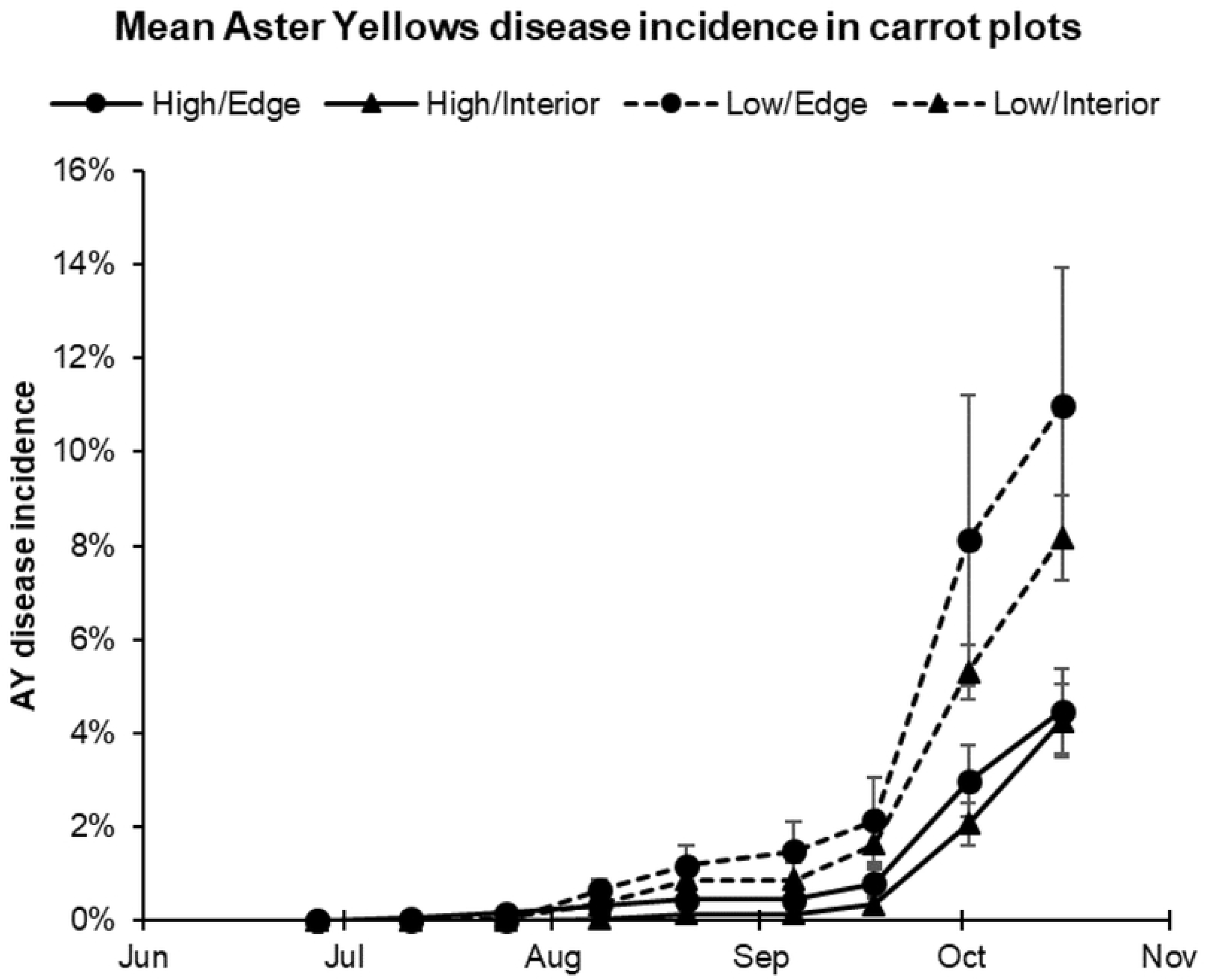
Mean Aster Yellows disease incidence in carrot plantings by planting density (high/low) and plot location within the field (edge/interior).

Aster Yellows disease incidence appeared to be influenced by planting density (high versus low density), by plot location within the field (field-edge versus interior plots), and was strongly dependent on date. To evaluate each of these factors, a mixed model was constructed to include *density* (high/low), *plot location* (edge/interior), and *week number*, and all two-way interactions as fixed effects, and *plot* as a random effect (**Supplemental Table S4**). In the incidence model, *density* (F=9.06, P=0.0083), *week* (F=479.49, P<0.0001), and the *density*week* interaction (F=26.81, P<0.0001) were significant. The random effect of plot was also significant (Wald’s p-value=0.040), indicating disease incidence differences between specific plots in the field independent of all other fixed effects. Plot location within the field showed a non-significant trend (F=2.63, p=0.12). This model confirms a significant difference in disease incidence between the high-density and low-density plantings, as well as a difference in the rate of change in disease progress between the two densities. However, a similar mixed model using only the number of diseased carrots per plot rather than the stand count-adjusted disease incidence values only, showed week as a significant component (f=497.82, p<.0001), with density, location, and all interaction terms non-significant. This suggests that the progression of AY through both the high- and low-density plantings was not limited by the number of available host plants, but rather by other factors.

### Aster leafhopper abundance and infectivity

Cumulative ALH abundance and rates of AYp-carrying ALH peaked in late August of 2018 and followed a similar bell-shaped distribution over both high and low planting densities. As the principal insect vector of the AYp in Wisconsin, ALH populations were monitored to evaluate vector pressure throughout the 2018 growing season in the high- and low-density carrot plots. From each leafhopper collection, a subset of adult ALH was assayed for phytoplasma to quantify the infectivity of the in-field leafhopper populations at unique time points throughout the study. Cumulative ALH abundance and rates of AYp-carrying ALH captured within the two plots peaked in late August of 2018 and followed similar bell-shaped distributions over both high and low planting densities (**Figure 2**). Aster leafhopper abundance in the field peaked from late July through late August, with populations declining through September and October (**Figure 2**). The fraction of captured ALH carrying the AYp followed a similar pattern, with a mean peak infectivity of 8.75% coincident with peak ALH populations (**Figure 2**).

**Figure 2.**
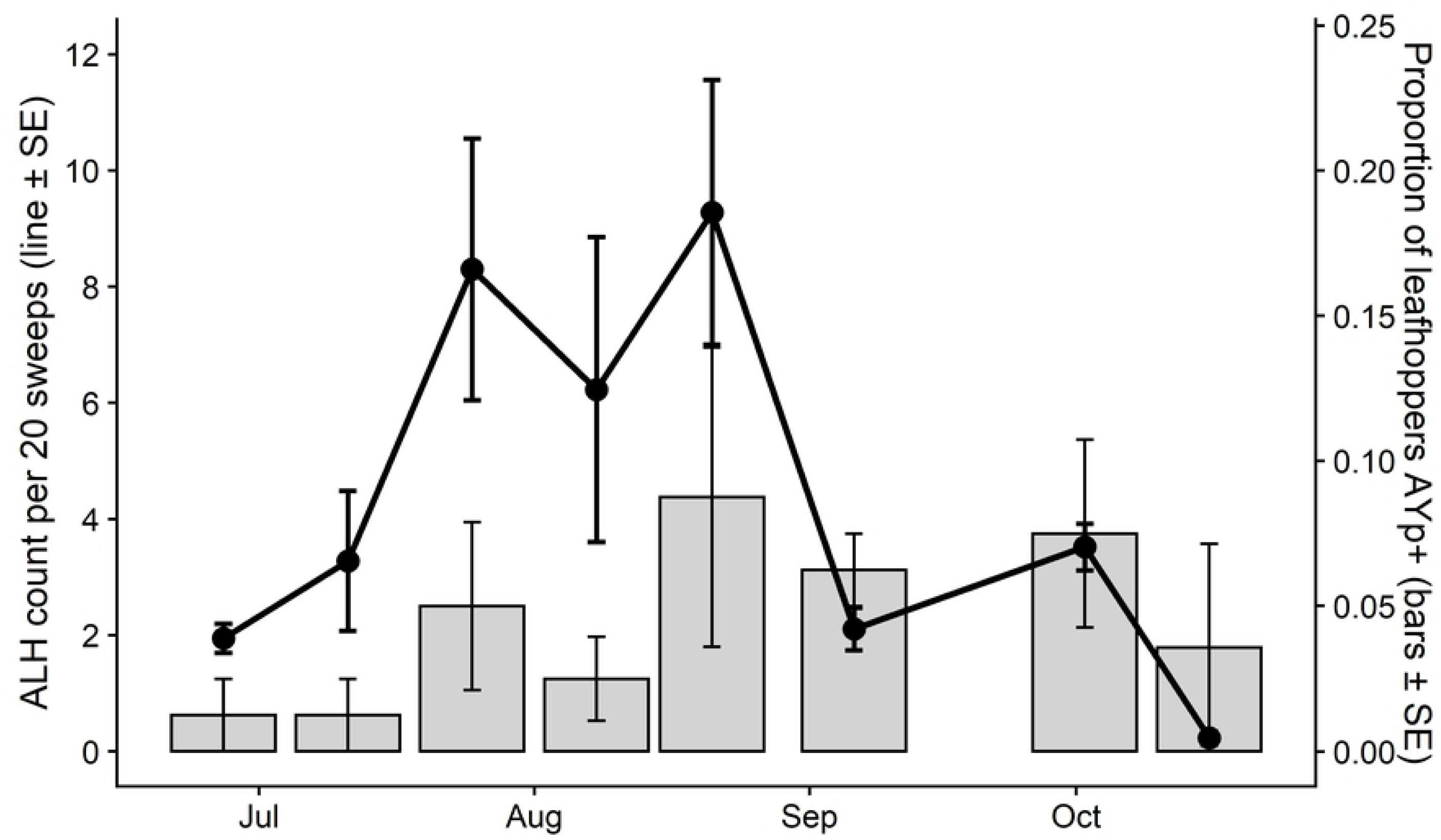
Mean abundance of aster leafhoppers and proportion of captured leafhoppers infected with AYp averaging over both planting densities within the carrot field.

Aster leafhopper abundance was greater in the high-density carrot plantings in July and August. Abundance peaked in the high-density planting at 32.85 ALH/ 20 sweeps. Further, phytoplasma-carrying ALH similarly peaked at 17.5% within the ALH populations in high density plantings on August 21 (**Figure 3**). The combination of high ALH counts together with high phytoplasma detections within the ALH coincide with high AYp disease incidence in carrot fields.

**Figure 3.**
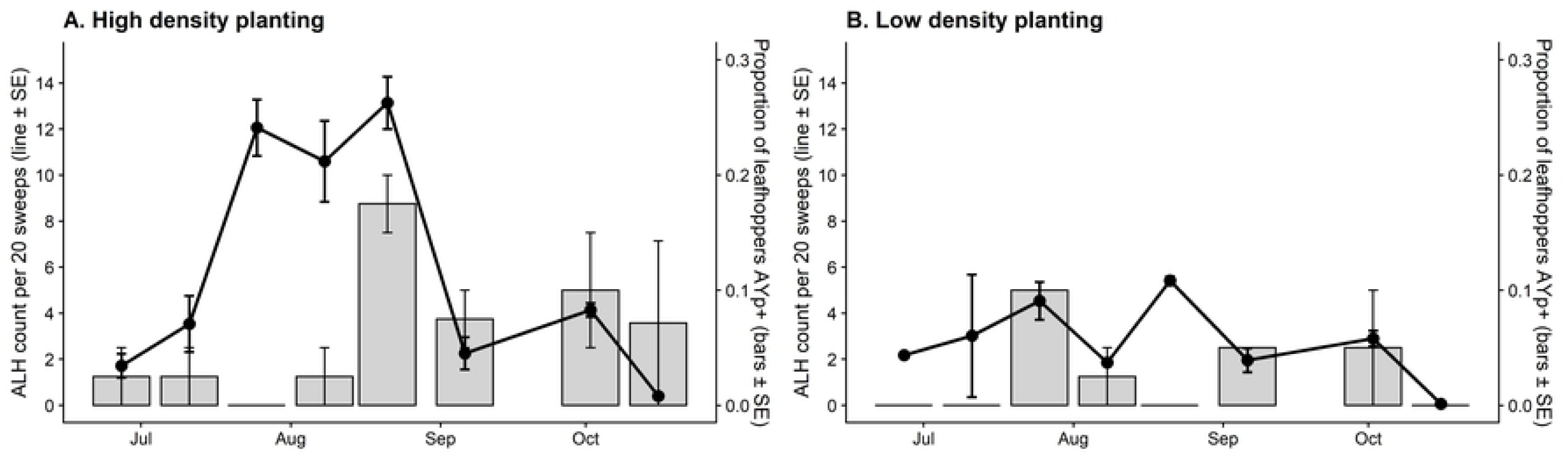
Abundance of aster leafhoppers and proportion of captured leafhoppers infected with AYp illustrating both high density (**A**) and low density (**B**) plantings in the carrot field.

### Aster yellows phytoplasma genotypic determination

Aster Yellows disease in carrots predominantly manifested late in the growing season with a higher proportion of AYp subgroup 16SrI-A (67%). In addition to evaluating carrot plots for disease progress, tissue samples from each symptomatic carrot (laboratory-confirmed AYp-positive samples) were genotyped to determine phytoplasma subgroup. In this experiment, only 16SrI-A and 16SrI-B AYp subgroups were observed within the field, with the overall subgroup composition changing significantly during the growing season (**Figure 4**). In the early portions of the sampling season, AYp samples were comprised of only 16SrI-B subgroup whereas a greater proportion of mid-season samples were mixed, containing varying proportions of both 16SrI-A and16SrI-B. The majority of AY diseased carrots possessed overt symptoms late in the growing season, and the greatest proportion of these infected carrots were classified as AYp subgroup 16SrI-A (67%)

**Figure 4.**
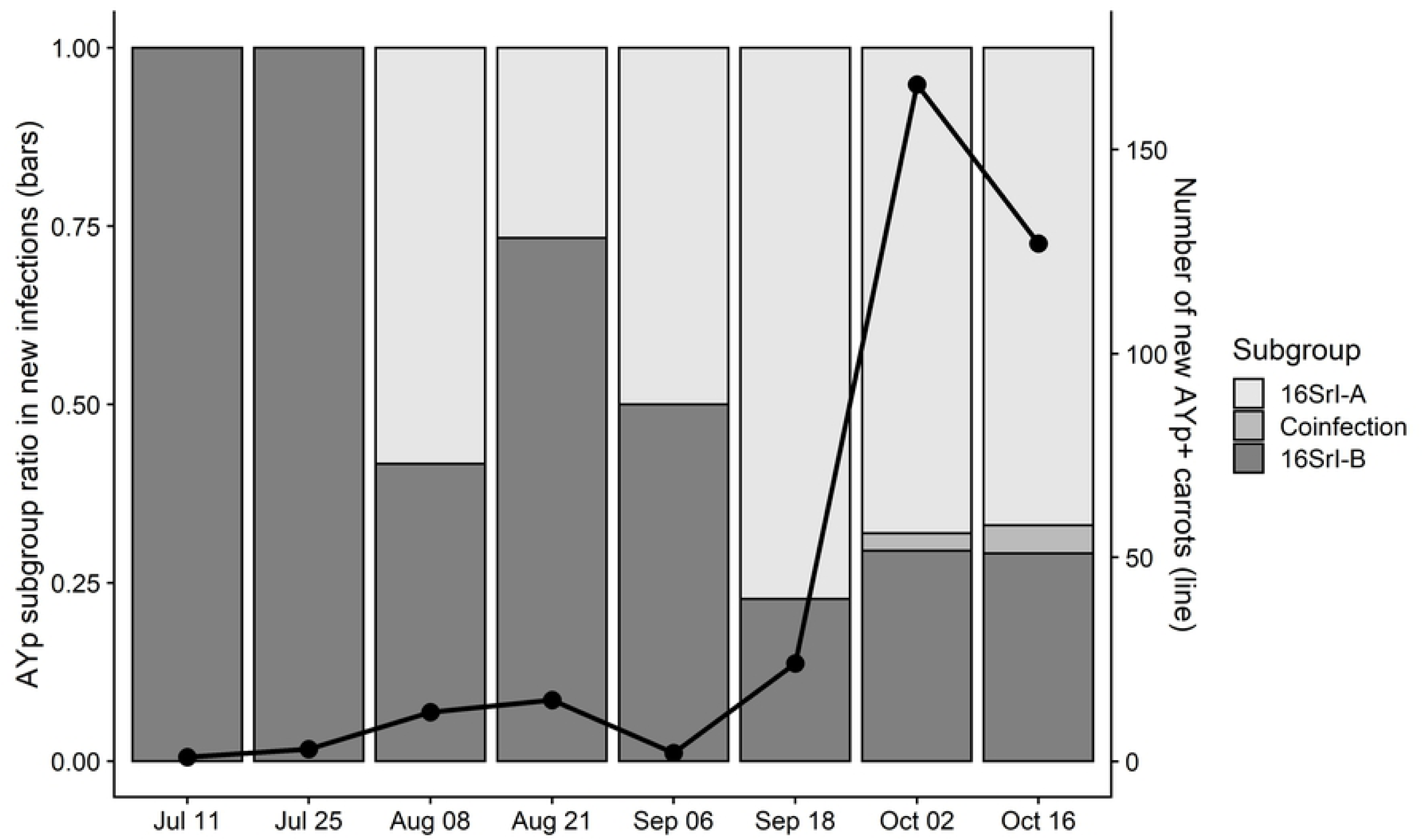
Number of new Aster Yellows phytoplasma infected carrots identified and sampled by date, illustrating the ratio of both AYp 16SrI-A and 16SrI-B subgroup proportions for each sample date.

### Secreted AY-WB effector identification

The distribution and proportion of SAP effectors depends upon AY subgroup designation. By employing ‘MonsterPlex’ parallel PCR amplification techniques, we were able to quantify the abundance of known effector sequences in the AYp genome and PMUs in 290 unique AYp samples collected from symptomatic carrots. To generate a “read ratio” for each effector, the number of reads per effector was normalized to the number of 16 rRNA gene reads for each sample. These normalized read values revealed significant differences in the presence of effectors genes between the two AYp subgroups observed in the investigation (**Figure 5**). From among the 20 effectors amplified, SAP11, 13, 15, 19, 45, and 68 were found to be specific to the 16SrI-A subgroup and not detected in the 16SrI-B subgroup (above an established threshold). Effectors SAP05, 06, 27, 35, 41, 42, 44, 48, 49, and 66 were present in both subgroups. Of the effectors present in both subgroups, SAP05, 06, 41, 42, and 66 amplifications generated more than double the amount of reads from samples that were infected with the 16SrI-A subgroup compared to the 16SrI-B subgroup. Both SAP06 and SAP66 were observed at an average read number 35.3 and 21.4 times greater, respectively, in the 16SrI-A subgroup compared to the 16SrI-B subgroup. The effectors with the highest number of reads in subgroup 16SrI-A samples were SAP41 (81.40 reads per 16 rRNA read) and SAP42 (78.34 reads per 16 rRNA read), while SAP48 (62.69 reads per 16 rRNA read) and SAP49 (68.34 reads per 16 rRNA read) had the highest copy within subgroup 16SrI-B (**Supplemental Table S2**). Only SAP44 generated slightly more reads from 16SrI-B versus 16SrI-A within infected samples.

**Figure 5.**
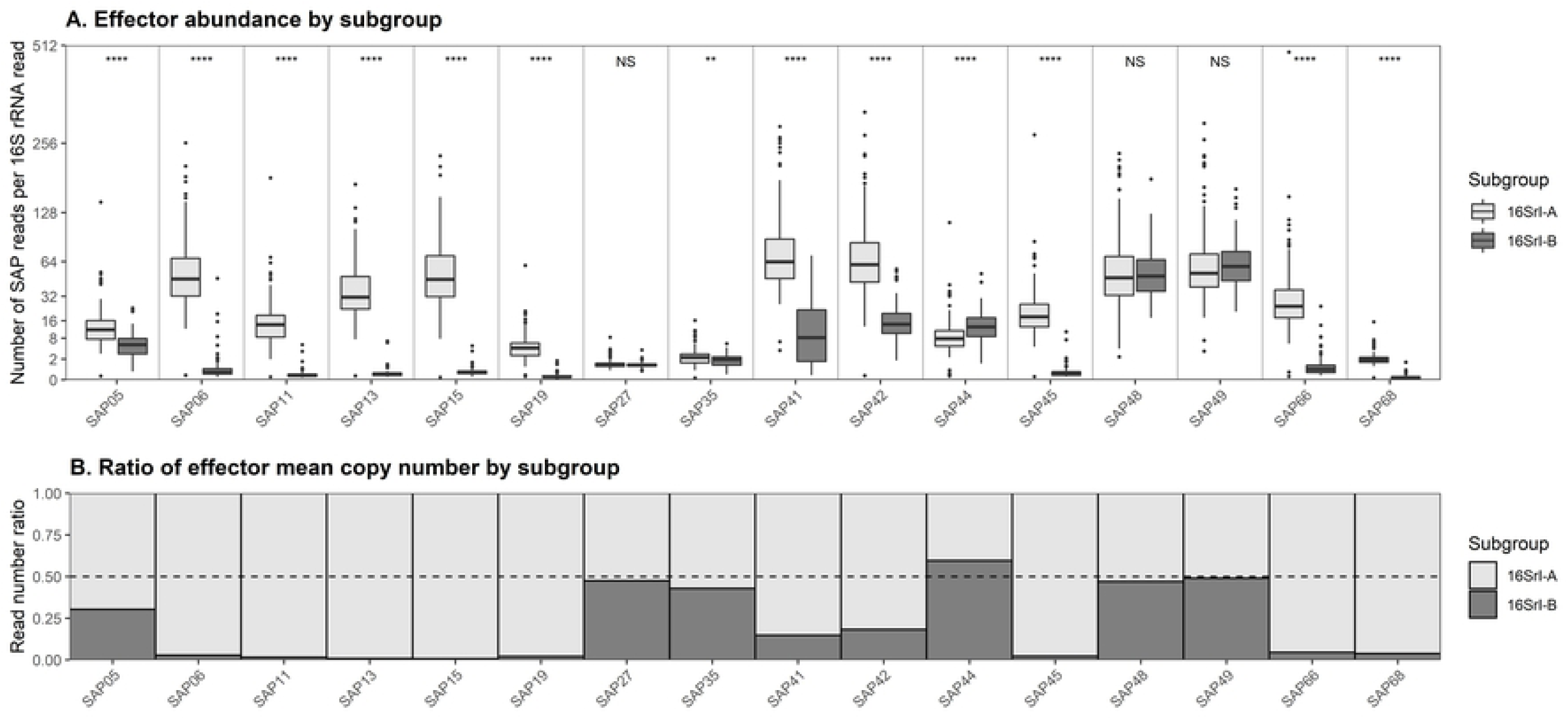
A) Effector read number within 16SrI-A and 16SrI-B subgroups. B) Effector ratio between 16SrI-A and 16SrI-B subgroups. Result of Chi-squared tests of significance comparing 16r RNA-normalized effector copy numbers shown along top of panel A (**** (P<0.0001), ** (P<0.01), NS=non-significant).

## Discussion and Conclusions

The movement and spread of AYp relies on multiple insect vectors, including the ALH [3]. Within susceptible agricultural crops the disease is monitored by growers and proactively controlled with insecticides when the vector is deemed in sufficiently high numbers [3]. However, if not controlled properly, AYp can manifest itself as AY disease, which can have significant ramifications in terms of crop yield and raw product quality. We sought to understand the progression of this disease in an insecticide-free environment to determine the ecological factors that influence AY disease progression. We examined the variables: time, planting density, location within the field (edge versus interior plot effects) and the associated genetics of the AYp (subgroup designation and effector composition) and hypothesized that disease progression within carrot fields by AYp was not random. However, we determined that the progression of AY disease is influenced by more than just the genotypic construct of AYp; it is also influenced by sample location, planting density and time during the crop season.

The overall progression of AY disease was influenced by sample location within field, planting density, and time of year. We observed that AY disease progressed at higher rates at lower planting densities. This suggests that infected leafhoppers were perhaps more readily attracted to infected plants, or that infected plants had greater apparency to mobile insects in less dense aggregations. Further, plants located along the outside edges of sample plots had a higher likelihood of infection by AYp, though only a non-significant trend (P=0.12). This could suggest that the movement of leafhoppers is directional and that leafhoppers are colonizing the field from the plot edges and inward. Similar to other members of the Hemiptera, adult leafhopper host location cues involve contrasts, increasing the likelihood that landing and initial inoculations may occur along field or plot edges. Leafhoppers were observed within our field as early as June 21, 2018, with the highest abundance on August 21, 2018. The highest rates of new infections were observed several weeks after peak leafhopper counts, in mid-September, which roughly corresponds to the previously described period of latency for development of visual AY symptoms in newly infected carrots, which can vary from 2-3 weeks [29]. While the observed infection rates within the low-density carrots were statistically higher than the high-density planting, the actual, and non-adjusted numbers of carrots infected in the low- and high-density planting were statistically similar. Although overall ALH numbers and the associated incidence of AYp infection within the insects was greater in the high-density plantings, the incidence of AYp infection within susceptible carrots (carrot cultivars that have been evaluated to be susceptible to infection by the AYp, and develop overt symptoms as a result of AYp infection) [30] was correspondingly lower, suggesting that the numbers of potentially inoculative insect vectors may not be limiting disease progress.

It has been previously established that AYp consists of multiple, genetically-distinct subgroups that can cause AY disease in carrots [11]. In Wisconsin there are at least two AYp subgroups, 16SrI-A and 16SrI-B, that are known AY disease agents [2, 11]. These earlier observations correspond to the findings of this study where we observed 16SrI-A, 16SrI-B, or a coinfection with both subgroups present in our experimental fields. The predominant subgroup at the beginning of the growing season was represented by 16SrI-B (100%), however as the season progressed the subgroup composition significantly shifted to 16SrI-A (67%) by the end of the season. The higher proportion of 16SrI-B at the beginning of the season could be an artifact of the initial low number of infected carrots. However, the shift in subgroup proportion is important to highlight and suggests that the 16SrI-A subgroup is being selected for in comparison to the 16SrI-B subgroup within our experiment. We did detect a small fraction of carrots infected with both 16SrI-A and 16SrI-B, suggesting that the pathogen can co-occur within the same host. Differences in SAP effector genes between the 16SrI-A and 16SrI-B subgroups may contribute to the higher abundance of 16SrI-A later in the season.

Secreted AY-WB effector genes were detected in symptomatic carrots in Wisconsin. Secreted AY-WB genes were amplified from field-collected plants infected with 16SrI-A phytoplasmas, consistent with primers designed to the 16SrI-A AY-WB phytoplasma genome. In addition, some effector genes were present in field-collected 16SrI-B infected plants, indicating that SAPs are present in both the 16SrI-A and 16SrI-B genomes found in Wisconsin. Differences in effector repertoires between the 16SrI-A and 16SrI-B subgroups could contribute to the distinct disease progression dynamics that are observed in the fields of Wisconsin. Secreted AY-WB proteins have been documented to manipulate host presentation to insect vectors. The effectors have been shown to have important biological ramifications on insect vector colonization (attractiveness) and reproduction (fitness). The exact function of all effectors investigated is still unknown. However, the functions of SAP11 and SAP54 have been well documented. Secreted AY-WB protein11 promotes oviposition (egg laying) by gravid, adult female ALH and SAP54 attracts insect vectors to plants by suppressing innate immune responses to herbivores. We noted SAP11 effectors were significantly more abundant in 16SrI-A. It is possible that the SAP11 effector may be more effective at promoting 16SrI-A phytoplasmas when there are more leafhoppers. However, how SAP11 modulates phytoplasma dynamics in a field situation remains to be determined. Overall, the effector proportion was significantly skewed towards 16SrI-A. Only SAP44 was significantly more abundant in the 16SrI-B phytoplasma subgroup. This observation suggests that subgroup 16SrI-A could be the predominant subgroup found in our field location due to the enhanced potential for pathogen spread associated with the increased proportion of effectors within this subgroup.

The data presented here represent an evaluation of a select set of factors which may influence progression of AY disease in susceptible carrots within central Wisconsin. We examined the factors: time during the growing season, initial planting density, sample location within the field and AYp subgroup and effector composition. Other factors could be evaluated in future studies to further complement our current understanding of the AY disease system including temperature, latitude, cultivar and cropping system. Here we demonstrate that AY disease was greater along plot edges and adjusted, final season incidence was greater in plots with lower, initial planting density. We also examined the genetic makeup of the phytoplasma and correlated subgroup 16SrI-A to higher effector proportion and greater disease spread. This information will lead to a better understanding of AYp movement in commercial crops, and contribute new knowledge towards describing factors that contribute to disease progress in susceptible carrots.

## Acknowledgements

The authors would like to acknowledge support from the Wisconsin Potato and Vegetable Growers Association and the associated agricultural community. We would also like to acknowledge Floodlight Genomics for guidance with their MonsterPlex Technology. We thank the farm staff at the Hancock Agricultural Research Station for planting and managing the plots throughout the season. This research was supported by Human Frontiers Science Program grant (HFSP RGP0024/2015) with additional funding from the Plant Health Institute Strategy Programme (BB/P012574/1) and the John Innes Foundation.

## Supplemental material

**Supplemental Table S1:** Primers

**Supplemental Table S2:** Effector abundance by subgroup

**Supplemental Table S3:** Carrot stand counts

**Supplemental Table S4:** Disease progression mixed model results

